# Are executive function and neuroanatomy in ADHD modulated by bilingualism?

**DOI:** 10.64898/2026.05.13.724877

**Authors:** Anushka Oak, Iria S. Gutierrez-Schieferl, Guinevere F. Eden

**Author notes:** **Corresponding Author** Guinevere Eden, D.Phil., Department of Pediatrics, Georgetown University Medical Center, Suite 150, Building D, 4000 Reservoir Road, NW, Washington, DC 20057, USA. **Co-author Details** Anushka Oak, address as above and, Iria Gutierrez-Schieferl, address as above and.

## Abstract

It has been proposed that bilinguals have better executive function (EF) arising from the constant selection of one language while inhibiting the other, and gray matter has been found to differ in bilinguals in regions linked to EF (frontal-parietal and subcortical structures). Attention Deficit Hyperactivity Disorder (ADHD) is associated with poorer EF and neuroanatomical differences underlying EF. Given the EF ‘advantage’ in bilinguals, we investigated whether a bilingual experience affects EF performance and brain structure differentially in those with ADHD. Using the Adolescent Brain and Cognitive Development Study, we compared early Spanish-English bilinguals and English-speaking monolinguals with and without ADHD. ANOVAs for the Flanker, Working Memory, and Card Sort Tasks revealed no main effects of Language Experience (Bilingual versus Monolingual), a main effect of Diagnostic Group for Card Sort (ADHD worse than Controls), and no interaction effects on performance for any task. ANOVAs for gray matter volume (GMV) revealed a main effect of Language Experience in many regions, a main effect of Diagnostic Group in some regions, but no interactions. GMV in left thalamus was affected by both ADHD and bilingualism, but the effect of ADHD was not significantly diminished or enhanced by the dual-language experience. For cortical thickness, there was a main effect of Language Experience in several regions, no main effect of Diagnostic Group, and no interactions. Taken together, bilingualism has some impact on EF performance, a strong impact on neuroanatomy, but there was no disproportionate impact by bilingualism on the differences caused by ADHD for any measure.

**Research Highlights:** Executive function and brain structure differ in ADHD and in bilinguals, prompting the need to investigate interactive effects.

Bilingualism did not disproportionately affect performance differences in ADHD for executive function, nor for gray matter volume or for cortical thickness differences in ADHD.

Gray matter volume was less in ADHD than non-ADHD, as well as greater in bilinguals than monolinguals in the left thalamus, but without interaction effect.

These independent effects indicate that the brain basis of ADHD is not impacted by a dual-language experience.

## Introduction

Both bilingualism and attention-deficit/hyperactivity disorder (ADHD) have been independently linked to differences in executive function (EF). EF is a constellation of top-down mental processes including inhibitory control, working memory, and cognitive flexibility that support goal-directed behavior (Diamond, 2013; Miyake et al., 2000) and are critical to children’s academic success (Horowitz-Kraus et al., 2026). In the field of bilingualism, it has been suggested that the constant selection of one language while inhibiting the other (Green, 1998; Green & Abutalebi, 2013) results in *better* EF in bilinguals (Bialystok et al., 2009). On the other hand, the developmental disorder ADHD, characterized by inattentive, hyperactive, and/or impulsive symptoms, is associated with *poorer* EF. The purported strength of EF in bilinguals raises the intriguing question of whether bilingualism plays a counteractive role on ADHD EF deficits, making the study of bilingualism, ADHD and their intersection a compelling, yet underexplored area of inquiry(Köder et al., 2022). There are strong reasons to study bilinguals with ADHD, given that bilinguals make up over 50% of the world population (Grosjean, 2021) and ADHD affects approximately 5-10% of children (Y. Li et al., 2023; Polanczyk et al., 2015). Bilingualism and ADHD are each associated with alterations of frontal-parietal and subcortical structures, yet to date neuroanatomical studies of ADHD have not taken bilingualism into consideration. Importantly, if the brain basis of ADHD is different in bilinguals than monolinguals, future studies of ADHD would need to account for this by distinguishing between bilingual and monolingual participants, and there could be translational implications for the diagnosis and treatment of bilinguals with ADHD.

Here, we address the question of whether EF task performance and brain structure manifest differently in those with ADHD who are bilingual relative to those with ADHD who are monolingual. Evidence that bilinguals have better EF than monolinguals on tasks involving inhibitory control (inference control, response inhibition), working memory, and cognitive flexibility (Bialystok & Martin, 2004; Carlson & Meltzoff, 2008; Morales et al., 2013) raises the possibility that these may counteract the EF deficits brought on by ADHD in bilinguals with ADHD. Poor EF has been central to theories of ADHD (Barkley, 2006; Castellanos et al., 2006; Rubia et al., 2018) and studies have reported poor inhibitory control, especially interference control on the Flanker task (Mullane et al., 2009) and response inhibition on the Stop-Signal Task (Schachar et al., 2000), poor working memory (Martinussen et al., 2005), and cognitive flexibility impairments (Tsuchiya et al., 2005). However, it should also be noted that the findings in both fields have not gone unchallenged. Recent work (Dick et al., 2019) and meta-analyses (Lehtonen et al., 2018; Lowe et al., 2021; Paap, 2019) do not confirm better EF in bilinguals than monolinguals, and while EF deficits in ADHD are well documented, recent population-based studies have reported smaller effect sizes than those found in earlier convenience samples (Cordova et al., 2022; Elosúa et al., 2017; Willcutt et al., 2005). Nevertheless, given the overall findings it is important to investigate the intersection of bilingualism and ADHD and so far, the findings have been mixed. A review by Köder et al. (2022) described nine studies on the combined effects of bilingualism and ADHD on the performance of EF tasks (Bialystok et al., 2017; Chung-Fat-Yim et al., 2020; Goh et al., 2020; Hardy et al., 2021; Mor et al., 2015; Sharma et al., 2019; Sharma et al., 2022; Sorge et al., 2017; Toppelberg et al., 2002), and concluded that no consistent pattern emerged. Notably these studies often contained multiple measures of EF, but bilingualism neither reliably helped nor hurt the group with ADHD in their performance on the task within the same study, or within the same category of EF tasks across studies. Thus, evidence for an interaction between language experience and diagnostic group on EF tasks is weak. Here we address this question with multiple EF measures in a group of participants that is larger than many of these prior studies and, importantly, in conjunction with brain anatomy.

Neuroanatomical differences have been reported for bilingualism, and, independently, for ADHD. Gray matter volume (GMV) studies of bilinguals suggest neuroanatomical differences across cortical and subcortical regions relative to monolinguals, particularly more GMV in areas supporting executive function, such as prefrontal, anterior cingulate, parietal, and temporal cortices (García-Pentón et al., 2019; Olulade et al., 2016; Schug et al., 2022), as well as basal ganglia and thalamus (Burgaleta et al., 2016; Nguyen et al., 2024; Pliatsikas et al., 2017). Olulade et al. (2016) and Schug et al. (2022) interpreted their GMV differences as reflecting engagement of EF networks through dual-language management (Green & Abutalebi, 2013), consistent with the involvement of frontal control regions in both language and executive function (Miller & Cohen, 2001; Niendam et al., 2012). Cortical thickness (CT) studies in bilinguals are more limited and reveal thinner cortex in bilinguals relative to monolinguals in frontal (Klein et al., 2014) and widely distributed regions (Vaughn et al., 2021). Turning to GMV studies of ADHD, these have also reported differences in brain regions implicated in EF. Specifically, less GMV in ADHD in prefrontal, orbitofrontal and anterior cingulate cortices (Frodl & Skokauskas, 2012; Norman et al., 2016), as well as subcortical regions supporting EF (basal ganglia and thalamus (Greven et al., 2015; Hoogman et al., 2017; C. S. Li et al., 2022; Nakao et al., 2011)). CT studies of ADHD similarly report thinner cortex in frontal and parietal regions (Narr et al., 2009; Shaw et al., 2006; Silk et al., 2016). However, taking the neuroanatomical studies of bilingualism and of ADHD into consideration, not all have supported the notion of structural differences in these groups, as demonstrated by a meta-analysis report of low convergence of findings for GMV differences in bilinguals (Danylkiv & Krafnick, 2020); or null findings for cortical thickness differences in bilinguals (Nguyen et al., 2023) and in studies of ADHD (Bernanke et al., 2022; Sarabin et al., 2023).

Given the evidence on potential involvement of fronto-parietal and subcortical regions in bilingualism and in ADHD, there is a strong motivation for an investigation into whether bilingualism influences brain structure in ADHD. The central question is whether the bilingual experience goes beyond an additive effect (manifesting equally in those with and without ADHD) and exerts an interactive effect such that the neuroanatomical manifestation of ADHD is either softened or enhanced in bilinguals. An impact of bilingualism that reduces the effects observed in monolinguals with ADHD, would suggest a protective benefit of bilingualism. However, a counteractive effect of any kind brought about by the dual-language experience would be important for informing accurate brain models of ADHD, recognizing that the neural signature of ADHD, derived from studies of monolinguals, may not entirely transfer to bilinguals.

The current study focused on cultural early bilinguals of Spanish and English (acquisition around ages 6–7 prior to formal schooling), which are prevalent in the U.S. and represent a relatively homogeneous group of bilinguals given the similar manner by which they came to acquire both languages (at a young age and as part of their home environment). We used data from the Adolescent Brain Cognitive Development (ABCD) Study, which is demographically representative of the U.S. We measured GMV, the dominant neuroanatomical metric in studies of bilingualism and ADHD, and CT, given the transition to the use of this measure due to its specificity. We conducted a factorial analysis to test for a main effect of Language Experience (Bilingual vs. Monolingual Groups), main effect of Diagnostic Group (ADHD vs. Control Groups), and their interaction on EF performance and on brain structure, the interaction gauging the differential impact of a dual-language experience on ADHD. Based on prior work noting differences in EF and brain structure (fronto-parietal and subcortical regions) in bilinguals and in ADHD, that run in opposite directions, one would hypothesize an interaction effect. Independent of the outcome, our results will inform whether the bilingual experience has any modifying effect on ADHD.

## Methods

### Participants

We obtained data from the ABCD Study Annual Release 5.0, an ongoing study following 11,868 youths across 21 collection sites (Garavan et al., 2018) and selected for the groups as described below and in Supplementary Figure 1. Unless otherwise noted, data are from the baseline visit, at which time participants were 9-10 years old. First, participants with missing values in the criteria variables were removed. Of the 11,557 available participants, we included participants with non-verbal reasoning, above 3 (approximately equivalent to a standard score above 70, or the 2nd percentile) on the Matrix Reasoning subtest from the Wechsler Intelligence Test for Children-V (Wechsler, 2014) after which 11,402 participants remained. Similarly, we only included participants with verbal reasoning above 70 on the NIH Toolbox Picture Vocabulary Test (Gershon et al., 2013, 2014), after which 11,334 participants remained. Per DSM-5 criteria (American Psychiatric Association, 2013), ADHD psychosis can present as inattention and other ADHD symptoms (Cordova et al., 2022) and we therefore excluded children with a current diagnosis of Unspecified Schizophrenia Spectrum or Other Psychotic Disorder on the KSADS-COMP, leaving 11,301 participants.

### Diagnostic Groups: ADHD and non-ADHD Controls

We then assigned the remaining participants into ADHD and non-ADHD Control Groups based on parent-reported attention problems, measured using the Attention Problems syndrome scale of the Child Behavior Checklist (CBCL; Achenbach & Ruffle, 2000), which Cordova et al. (2022) identified as the most useful dimensional measure of ADHD in the ABCD dataset. Participants with a score of 65 and above were assigned as ADHD, resulting in 821 participants. This represents 7.1% of the sample, slightly lower than reported by Cordova et al., 2022 (8.54%), likely due to the exclusion of participants with lower verbal skills. Those scoring below 65 were designated as non-ADHD controls, resulting in 10,476 participants.

### Language Experience: Bilingual Groups

Next, we used the parent and youth demographic questionnaires from the ABCD Study to identify bilingual participants amongst these ADHD and non-ADHD Control Groups. We first selected those who responded ‘Yes’ to the question, ‘Besides English, do you speak or understand another language or dialect?’ on the Youth Acculturation Survey (YAS), modified from PhenX, which left 278 in the ADHD Group and 3,945 in the non-ADHD Control Group. We then included those who answered ‘Spanish’ to the question ‘What other language do you speak or understand besides English?’ on the YAS resulting in 204 in the ADHD Group and 2,751 in the non-ADHD Control Group. Next, we identified those who responded ‘Non-English Language all the time’, ‘Non-English Language most of the time’, or ‘Non-English Language and English equally’ when asked ‘What language do you speak with most of your family?’ on the YAS to focus on heritage speakers, resulting in 69 in the ADHD Group and 1,046 in the non-ADHD Control Group. Finally, we included children whose parents responded ‘Yes’ when asked ‘Do you consider the child Hispanic/Latino/Latina?’ on the Parent Demographics Survey. This yielded 61 cultural Spanish-English bilinguals in the ADHD Group and 970 cultural Spanish-English bilinguals in the non-ADHD Control Group.

### Language Experience: Monolingual Groups

We identified monolingual participants from those who met (821) or did not meet (10,476) the ADHD Group criteria. We included only those who had responded ‘No’ when asked, ‘Besides English, do you speak or understand another language or dialect?’ on the YAS which left 543 in the ADHD Group and 6,473 in the non-ADHD Control Group. We excluded those children whose parents reported the child’s native language to be something other than English on the Longitudinal Parent Demographics Survey at the One-Year Follow-Up time point (‘What is your child’s native language? In other words, what was the first language predominantly spoken to your child by their parent or guardian after birth?’), resulting in 495 in the ADHD Group and 6,114 in the non-ADHD Control Group. Further, we excluded children enrolled in a dual-language school (according to one-year follow-up data). These criteria ensured we included only those children with minimal if any exposure to languages other than English: 484 in the Monolingual ADHD Group and 6,003 in the Monolingual Control Group.

### Propensity Score Matching and Final Groups

After eliminating participants without T1-weighted MPRAGE structural brain images (according to the Image03 spreadsheet), participants were submitted to propensity score matching using the PsmPy package (Kline & Luo, 2022). To ensure any anatomical differences could not be attributed to the known associations with nonverbal reasoning (Schilling et al., 2013), verbal reasoning (Bialystok & Craik, 2010) or socioeconomic status (SES) (Brito et al., 2018; Lotze et al., 2020; Rakesh & Whittle, 2021), we matched the four groups on these variables using the Matrix Reasoning subtest from the Wechsler Intelligence Test for Children-V (Wechsler, 2014), the Picture Vocabulary Test (Gershon et al., 2013, 2014), and the Longitudinal Parent Demographics Survey used in the ABCD Study. SES was assessed using combined household income (Bins: 1= Less than $5,000; 2=$5,000 through $11,999; 3=$12,000 through $15,999; 4=$16,000 through $24,999; 5=$25,000 through $34,999; 6=$35,000 through $49,999; 7=$50,000 through $74,999; 8= $75,000 through $99,999; 9=$100,000 through $199,999; 10=$200,000 and greater). Propensity scores were calculated using logistic regression and groups were matched to the Bilingual ADHD Group (n =61) 1:1 without replacement, leading to groups of 61 participants in each of the four groups: Bilingual ADHD, Bilingual Control, Monolingual ADHD, and Monolingual Control Groups.

Following the first step of quality control of the MRI images (see below), several participants were excluded for the GMV analysis, leaving sample sizes of 59 (Bilingual ADHD), 57 (Bilingual Control), 55 (Monolingual ADHD), and 59 (Monolingual Control), but demographic (SES) and performance (nonverbal and verbal reasoning) measures remained matched between the groups. Following another step of quality control specific to the CT analysis (see below), three more participants were excluded, leaving sample sizes of 59 (Bilingual ADHD), 56 (Bilingual Control), 53 (Monolingual ADHD), and 59 (Monolingual Control), but demographic (SES) and performance (nonverbal and verbal reasoning) measures remained matched between the groups. Table 1 shows the participants (n=61 per group) prior to the exclusion based on MRI quality control, as these data were used in the study comparing EF performance.

**Table 1:**
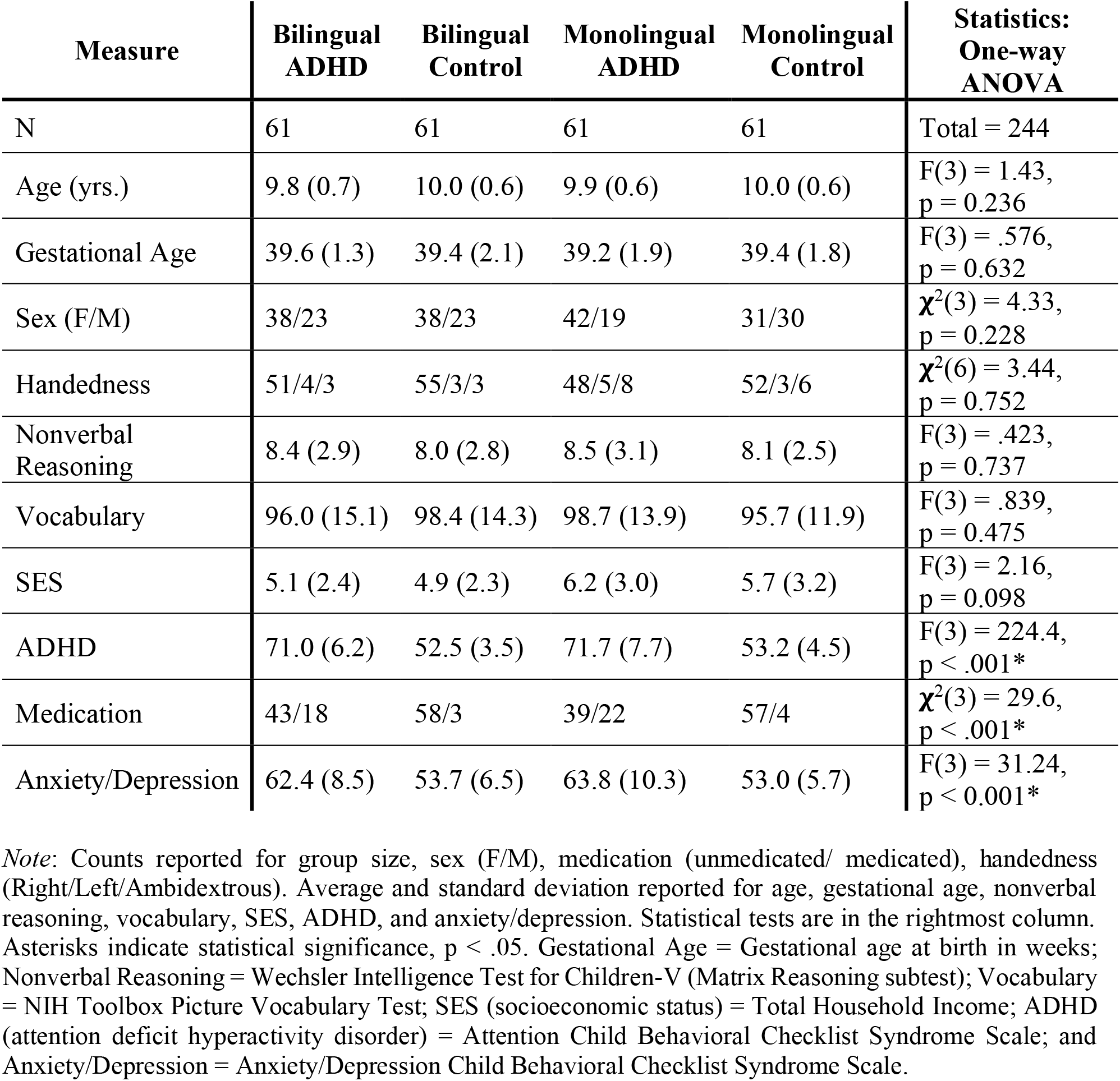
Participant Demographics for Early Bilingual and Monolingual ADHD and Controls.

### Behavioral Performance on Executive Function Tasks

We tested performance on three executive function tasks administered in the ABCD Study (Luciana et al., 2018). The Flanker task (Eriksen & Eriksen, 1974) measures interference control by assessing whether surrounding stimuli and the target are congruent or incongruent. The List Sort Working Memory task measures working memory by asking participants to sequence pictured items based on category and perceptual features (Tulsky et al., 2014). The Dimensional Change Card Sorting task (Zelazo, 2006) measures cognitive flexibility by requiring participants to sort objects by either shape or color, testing their ability to shift between rules. These tasks exemplify three of the core components of EF: inhibitory control, working memory and cognitive flexibility (Diamond, 2013; Miyake et al., 2000). We did not utilize the stop-signal task due to concerns about the interpretation of the results for this version of the test in the ABCD Study (Bissett et al., 2021).

### MRI Data Acquisition

Whole-brain T1-weighted MR images were acquired on 3T MR scanners with a 1 × 1 × 1 mm^3^ voxel resolution. Details on acquisition protocols can be found at https://abcdstudy.org/images/Protocol_Imaging_Sequences.pdf.

### MRI Data Quality Control

First, two researchers blinded to group membership assessed the MR images and excluded those images with obvious anatomical anomalies and/or a score of 1 or 2 on an image quality scale of 1 (severely distorted) to 5 (optimal). Twelve participants with image scores of 1 and 2 were eliminated. Before the CT analyses, there was an additional quality control step following image segmentation in CAT12, when all segmented images were assessed with the CAT12 Weighted Overall Image Quality. The resulting image quality rating (IQR) values [on a scale of 0.5 (excellent) to 10.5 (unacceptable)] were converted to percentage rating values based on the CAT12 manual (https://neuro-jena.github.io/cat12-help/). Images with ratings below 70% (i.e., IQR > 3.5) were excluded from the analyses. Also, the segmented gray matter images were visually inspected for skull-stripping and segmentation errors, and any images with obvious errors were excluded. These additional steps eliminated three additional participants.

### Executive Function Performance

We performed a 2 × 2 full factorial ANOVA on the four participant groups for accuracy on the Flanker Task, the List Sort Working Memory Task and the Dimensional Change Card Sorting Task. Statistical analyses were conducted using Jamovi (https://jamovi.org). Other performance and demographic variables were also compared using Jamovi.

### Gray Matter Volume

All images were preprocessed in SPM12 using the automated Voxel-Based Morphometry (VBM) (Ashburner & Friston, 2000). First, images were coregistered to the tissue probability map and then segmented into gray matter, white matter, and cerebrospinal fluid images. These images were then used to create a study-specific template and spatially normalized to the Montreal Neurological Institute (MNI) stereotaxic space via affine registration of the generated template to the MNI template using DARTEL (Ashburner, 2007). Next, images were smoothed with a Gaussian kernel of 8 mm full width at half-maximum. Finally, to reduce edge artifacts, the intensity threshold of the images was set to 0.2.

To test the main research question of differential effects of a dual-language experience in children with ADHD, we entered all four groups (Bilingual ADHD, Bilingual Control, Monolingual ADHD, and Monolingual Control) into a 2 × 2 full factorial analysis in SPM12 (voxel-wise height threshold of p < 0.005, uncorrected; cluster-level extent threshold of p < 0.05, FDR corrected) to test for the main effects of Diagnostic Group, main effect of Language Experience, and their interaction. Age in months, sex, pubertal stage, total GMV, socioeconomic status, study site, anxiety/depression on the CBCL syndrome scale, and stimulant medication were included as covariates of no interest. Stimulant medication information was derived from the ABCD parent demographic surveys. A binary variable was created, with 1 indicating use of one or more stimulant medications (e.g., Adderall, Ritalin, Vyvanse) during the previous two weeks and 0 indicating non-use (Owens et al., 2021).

### Cortical Thickness

Using the CAT12 (Gaser et al., 2022) toolbox in SPM12, a study-specific tissue probability map was created using the Template-O-Matic (TOM8) toolbox (Wilke et al., 2008), matched to the age and sex distribution of participants and used in the segmentation of the MR images. The images were then smoothed with a Gaussian kernel of 10 mm full width at half-maximum.

To test the main research question, we used the same factorial design as described above, this time for cortical thickness (vertex-wise height threshold of p < 0.005, uncorrected; cluster-level extent threshold of p < 0.05, FDR corrected) and entering the covariates listed above, except for total GMV.

For both GMV and CT, the publicly available label4MRI package was used to determine Brodmann’s areas (BA) and anatomical labels of coordinates outputted by SPM12 (https://github.com/yunshiuan/label4MRI). This package uses the Atlas of Brodmann’s areas and the Automated Anatomical Labeling Atlas (Tzourio-Mazoyer et al., 2002) respectively.

## Results

### Demographics and Behavioral Performance

Table 1 provides demographic and performance averages of the four selected groups (n=61 each): Bilingual ADHD, Bilingual Control, Monolingual ADHD, and Monolingual Control Groups. Chi-square tests and one-way ANOVAs revealed that the groups did not differ in age, gestational age, sex distribution, or handedness. As expected, given the propensity score matching, the four groups also did not differ on nonverbal reasoning, verbal reasoning, or SES. Further, the ADHD and non-ADHD Control Groups differed on ADHD symptomatology, stimulant medication use, and anxiety/depression, reflecting diagnostic criteria, pharmacological treatment, and known comorbidities, respectively. Importantly, these measures did not differ between Bilingual and Monolingual Groups. For community of descent, see Supplementary Table 1.

### Executive Function Performance

As shown in Table 2, the 2 × 2 ANOVAs revealed no main effects of Language Experience on the Flanker, List Sort Working Memory or Dimensional Change Card Sort Tasks. There was a main effect of Diagnostic Group on the Dimensional Change Card Sorting Task (ADHD < Control; F(1,240) = 4.42, p = .036), but not the other two tasks. There were no interaction effects between Diagnostic Group (ADHD vs Control) and Language Experience (Bilingual vs. Monolingual) on performance on any of the three EF tasks.

**Table 2:** Executive Control Task Performance for Early Bilingual and Monolingual ADHD and Controls.

| Measure | Bilingual ADHD | Bilingual Control | Monolingual ADHD | Monolingual Control | Statistics: $2 \times 2$ ANOVA |
| --- | --- | --- | --- | --- | --- |
| Flanker | 90.1 (15.4) | 92.6 (13.1) | 92.9 (15.0) | 92.8 (14.6) | Language: $F(1,240) = 0.660$, $p = 0.417$<br>ADHD: $F(1,240) = 0.373$, $p = 0.542$<br>Interaction: $F(1,240) = 0.475$, $p = 0.491$ |
| List Sort Working Memory | 92.2 (11.0) | 94.9 (14.7) | 95.1 (14.5) | 94.5 (17.9) | Language: $F(1,240) = 0.44$, $p = 0.150$<br>ADHD: $F(1,240) = 0.31$, $p = 0.576$<br>Interaction: $F(1,240) = 0.78$, $p = 0.350$ |
| Dimensional Change Card Sorting | 90.6 (13.5) | 94.2 (15.5) | 90.3 (14.0) | 94.3 (13.7) | Language: $F(1,240) = 0.0040$, $p = 0.950$<br>ADHD: $F(1,240) = 4.42$, $p = 0.036^*$<br>Interaction: $F(1,240) = 0.0082$, $p = 0.928$ |

Note: Average accuracy and standard deviation for the executive control tasks. Results from $2 \times 2$ factorial ANOVAs testing for Main Effects of Language Experience, Main Effects of Diagnostic Group, and their interaction on performance are presented in the rightmost column. Asterisks indicate statistical significance, $p < .05$ .

**Table 3:**
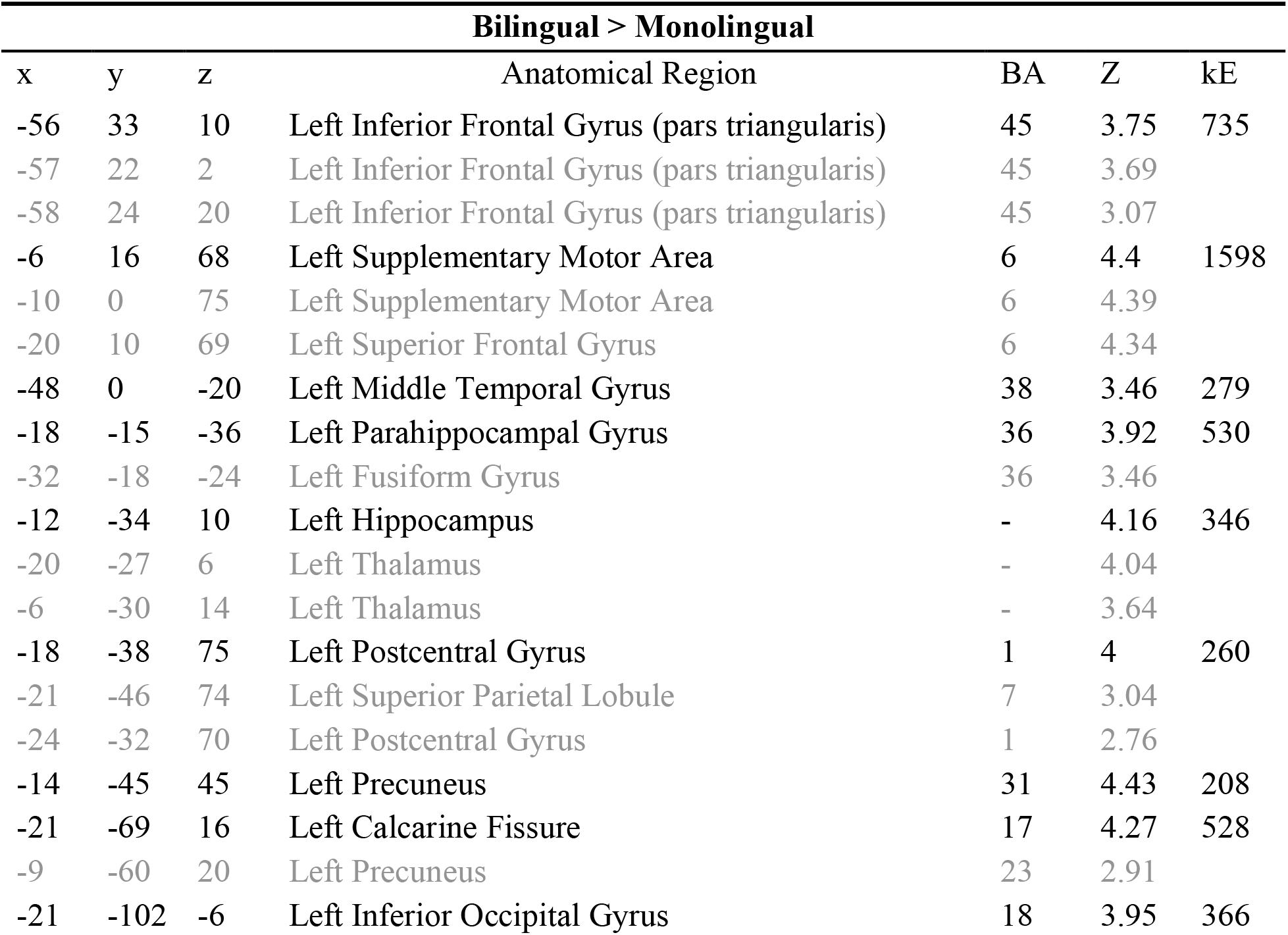
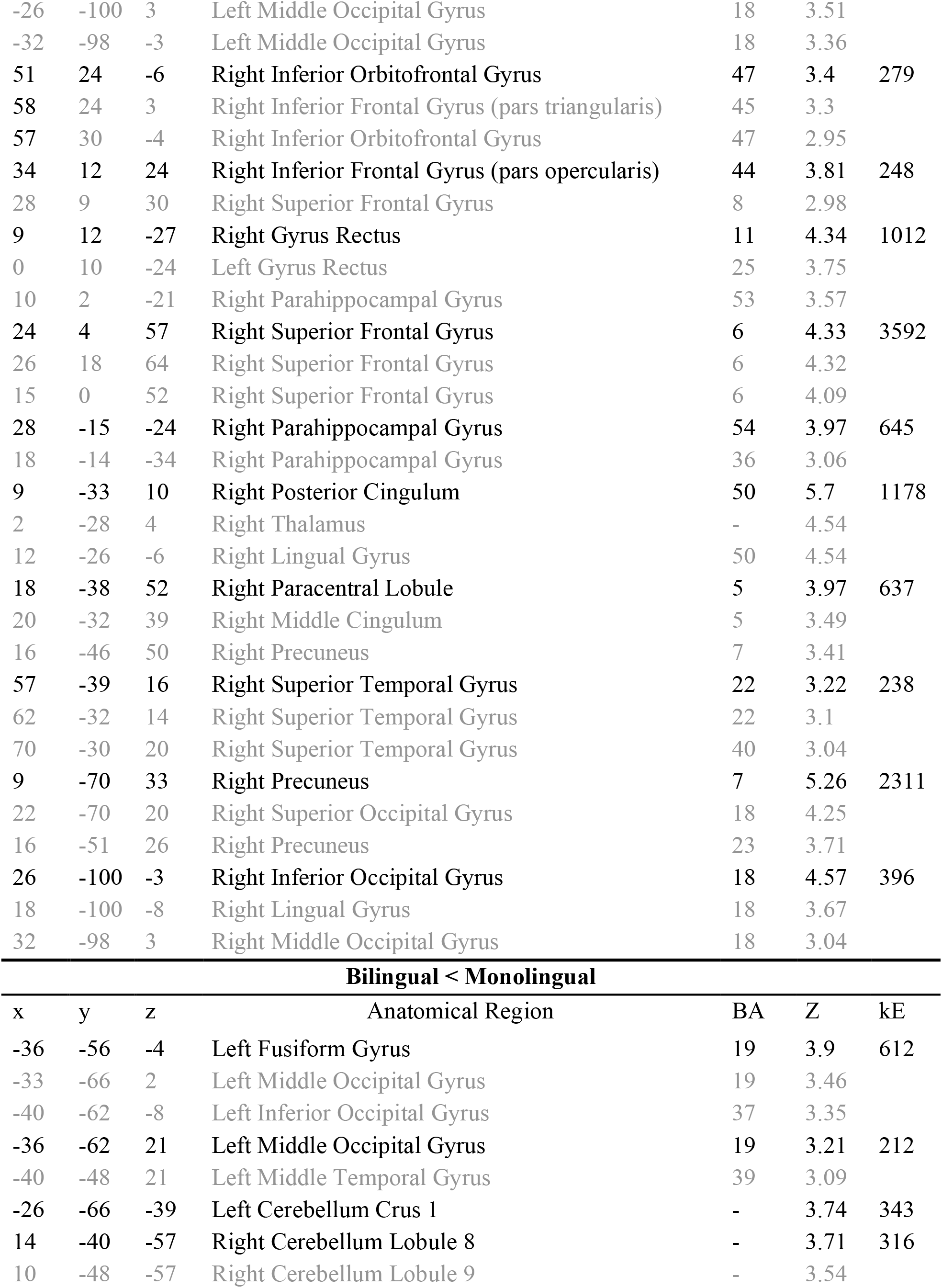
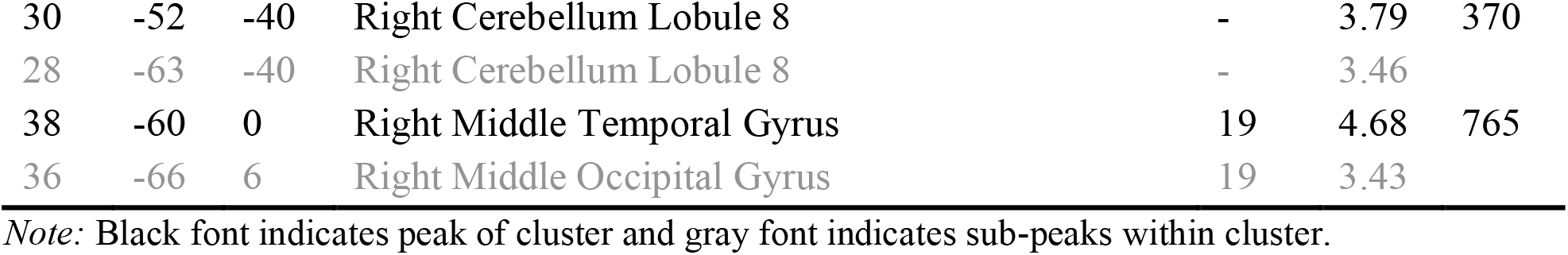
MNI Coordinates for Maxima of Main Effect of Language Experience on GMV.

**Table 4:**
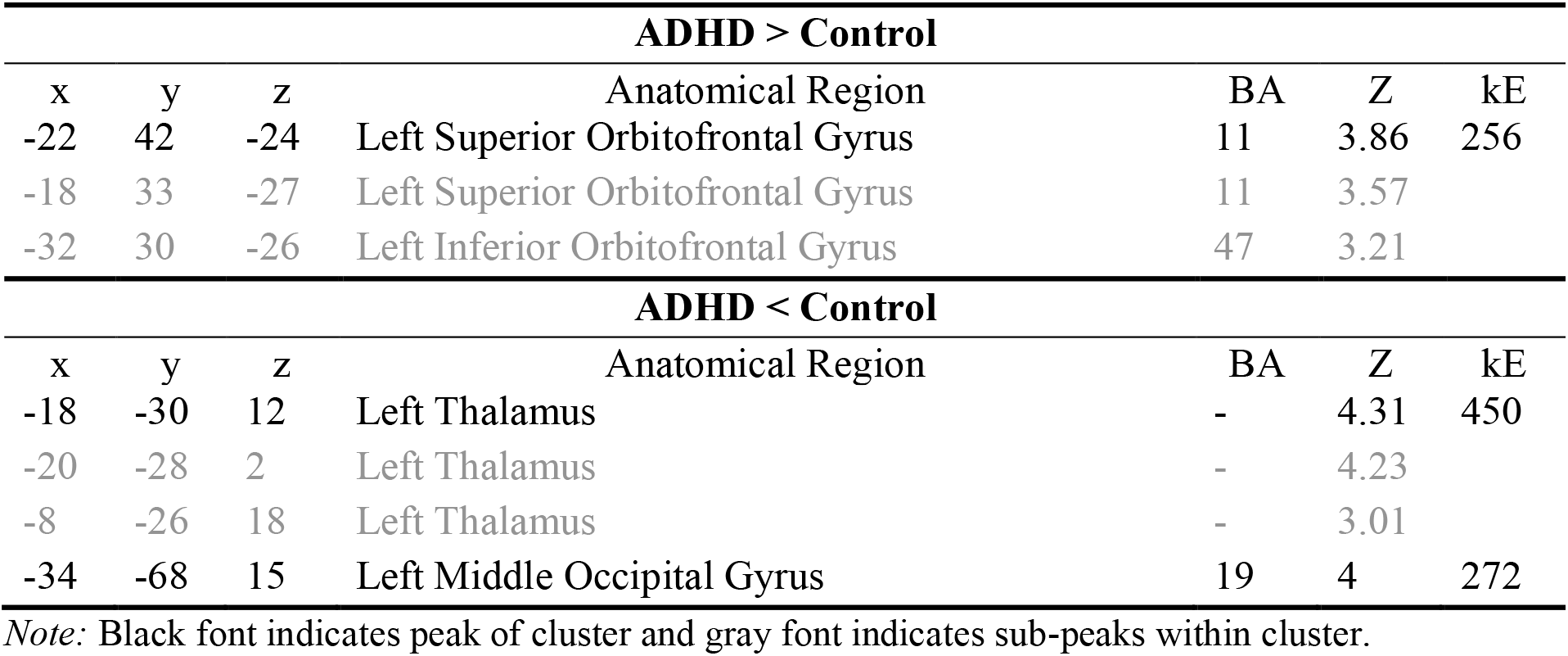
MNI Coordinates for Maxima of Main Effect of Diagnostic Group on GMV.

### Gray Matter Volume

#### GMV: Main Effect of Language Experience

As shown in Table 3 and Figure 1(A), group comparisons between bilinguals (Control and ADHD Groups combined) and monolinguals (Controls and ADHD Groups combined) showed widespread differences in gray matter volume (GMV). The main effect of Language Experience revealed 19 regions where the bilinguals had more GMV than the Monolinguals. Nine in the left hemisphere: (1) inferior frontal gyrus pars triangularis (BA 45), (2) supplementary motor area (BA 6) extending to the superior frontal gyrus (BA 6), (3) middle temporal gyrus (BA 38), (4) parahippocampal gyrus (BA 36) extending to the fusiform gyrus (BA 36), (5) hippocampus extending to the thalamus, (6) postcentral gyrus (BA 1) extending to the superior parietal lobule (BA 7), (7) precuneus (BA 31), (8) calcarine fissure (BA 17) extending to the precuneus (BA 23), and (9) inferior occipital gyrus (BA 18) extending to the middle occipital gyrus (BA 18). The remaining ten clusters had peaks in the right hemisphere: (1) inferior orbitofrontal gyrus (BA 47) extending to the inferior frontal gyrus pars triangularis (BA 45) and inferior orbitofrontal gyrus (BA 47), (2) inferior frontal gyrus pars opercularis (BA 44) extending to the superior frontal gyrus (BA 8), (3) gyrus rectus (BA 11) extending to the left gyrus rectus (BA 25) and parahippocampal gyrus (BA 53), (4) superior frontal gyrus (BA 6), (5) parahippocampal gyrus (BA 54) extending to the parahippocampal gyrus (BA 36), (6) posterior cingulum (BA 50) extending to the thalamus and lingual gyrus (BA 50), (7) paracentral lobule (BA 5) extending to the middle cingulum (BA 5) and precuneus (BA 7), (8) superior temporal gyrus (BA 22) extending to the superior temporal gyrus (BA 22 and BA 40), (9) precuneus (BA 7) extending to the superior occipital gyrus (BA 18) and precuneus (BA 23), and (10) inferior occipital gyrus (BA 18) extending to the lingual gyrus (BA 18) and middle occipital gyrus (BA 18). The opposite contrast revealed 6 regions where the bilinguals had less GMV than the Monolinguals. Three in the left hemisphere: (1) fusiform gyrus (BA 19) extending to the middle occipital gyrus (BA 19) and inferior occipital (BA 37), (2) middle occipital gyrus (BA 19) extending to the middle temporal gyrus (BA 39), and (3) cerebellum crus I (BA 19). The remaining three clusters had peaks in the right hemisphere where the second and third were in the same region: (1) cerebellum lobule VIII extending to cerebellum lobule IV, (2) cerebellum lobule VIII, and (3) middle temporal gyrus (BA 19) extending to the middle occipital gyrus (BA 19).

**Figure 1:**
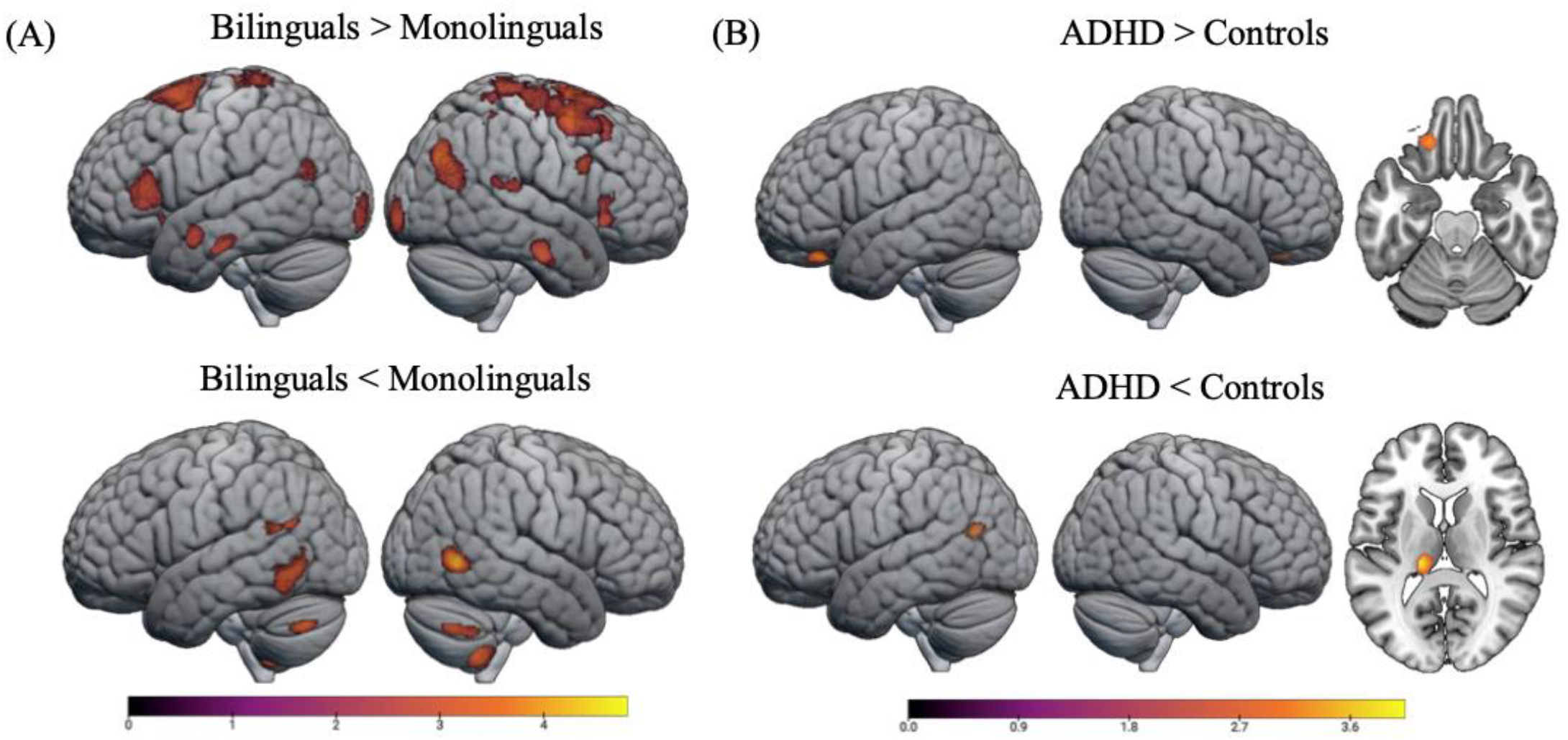
Gray Matter Volume Results. **(A)** Main Effect of Language Experience **(B)** Main Effect of Diagnostic Group on GMV. Voxel-wise height threshold p < 0.005, cluster-level extent threshold p < 0.05, FDR. Coordinates for significant clusters for the main effect of Language Experience are provided in Table 3, and for the main effect of Diagnostic Group in Table 4. (n = 230)

#### GMV: Main Effect of Diagnostic Group

As shown in Table 4 and Figure 1(B), the main effect of Diagnostic Group revealed one region where the ADHD Group had more GMV than the Control Group. This region was the left superior orbitofrontal gyrus (BA 11) extending to the inferior orbitofrontal gyrus (BA 47). The opposite contrast revealed two regions where the ADHD group (Monolinguals and Bilinguals Groups combined) had less GMV than the Control group (Monolinguals and Bilingual Groups combined): (1) left thalamus and (2) left middle occipital gyrus.

#### GMV: Diagnostic Group x Language Experience Interaction

There were no results from the interaction between Diagnostic Group and Language Experience on GMV.

### Cortical Thickness

#### CT: Main Effect of Language Experience

As shown in Table 5 and Figure 2, the main effect of Language Experience on CT revealed nine regions where the Bilingual Group (ADHD and Controls combined) had less CT than Monolinguals (ADHD and Controls combined). Six in the left hemisphere: (1) supplementary motor area (BA 8) extending to superior frontal gyrus and the supplementary motor area (BA 6), (2) middle frontal gyrus (BA 6) extending to the middle frontal gyrus (BA 8), (3) superior temporal pole (BA 38) extending to the middle temporal pole (BA 38), (4) precentral gyrus (BA 6) extending to the inferior frontal gyrus pars opercularis (BA 44) and postcentral gyrus (BA 4), (5) fusiform gyrus (BA 36), and (6) angular gyrus (BA 39) extending to the middle temporal gyrus (BA 21). The remaining three clusters were in the right hemisphere: (1) superior temporal pole (BA 38), (2) supplementary motor area (BA 6), and (3) precuneus (BA 31). The opposite contrast revealed no regions where the Bilingual Group had more CT than the Monolingual Group.

**Table 5:** MNI Coordinates for Maxima of Main Effect of Language Experience on CT.

| **Bilingual < Monolingual** |  |  |  |  |  |  |
| --- | --- | --- | --- | --- | --- | --- |
| x | y | z | Anatomical Region | BA | Z | kE |
| -6 | 20 | 55 | Left Supplementary Motor Area | 8 | 4.55 | 486 |
| -10 | 42 | 49 | Left Superior Frontal Gyrus | 8 | 3.76 |  |
| -8 | 4 | 69 | Left Supplementary Motor Area | 6 | 3.45 |  |
| -30 | 11 | 51 | Left Middle Frontal Gyrus | 6 | 4.55 | 195 |
| -23 | 19 | 44 | Left Middle Frontal Gyrus | 8 | 3.14 |  |
| -49 | 8 | -15 | Left Superior Temporal Pole | 38 | 4.74 | 223 |
| -39 | 17 | -36 | Left Middle Temporal Pole | 38 | 3.15 |  |
| -60 | 1 | 25 | Left Precentral Gyrus | 6 | 3.14 | 304 |
| -51 | 15 | 24 | Left Inferior Frontal Gyrus (pars opercularis) | 44 | 3.13 |  |
| -58 | -10 | 23 | Left Postcentral Gyrus | 4 | 3.09 |  |
| -31 | -25 | -26 | Left Fusiform Gyrus | 36 | 3.89 | 189 |
| -28 | -19 | -32 | Left Fusiform Gyrus | 36 | 3.81 |  |
| -30 | -11 | -33 | Left Fusiform Gyrus | 36 | 3.65 |  |
| -46 | -58 | 25 | Left Angular Gyrus | 39 | 4.86 | 190 |
| -46 | -50 | 11 | Left Middle Temporal Gyrus | 21 | 2.76 |  |
| 52 | 10 | -16 | Right Superior Temporal Pole | 38 | 5.03 | 160 |
| 5 | -14 | 57 | Right Supplementary Motor Area | 6 | 4.36 | 542 |
| 7 | 6 | 50 | Right Supplementary Motor Area | 6 | 3.74 |  |
| 14 | 11 | 62 | Right Supplementary Motor Area | 6 | 3.62 |  |
| 14 | -50 | 36 | Right Precuneus | 31 | 4.08 | 150 |
| 3 | -50 | 27 | Right Precuneus | 23 | 2.9 |  |
| 5 | -55 | 18 | Right Precuneus | 23 | 2.62 |  |
| **Bilingual > Monolingual** |  |  |  |  |  |  |
| None |  |  |  |  |  |  |

*Note:* Black font indicates peak of cluster and gray font indicates sub-peaks within cluster.
*CT: Main Effect of Diagnostic Group*

**Figure 2:**
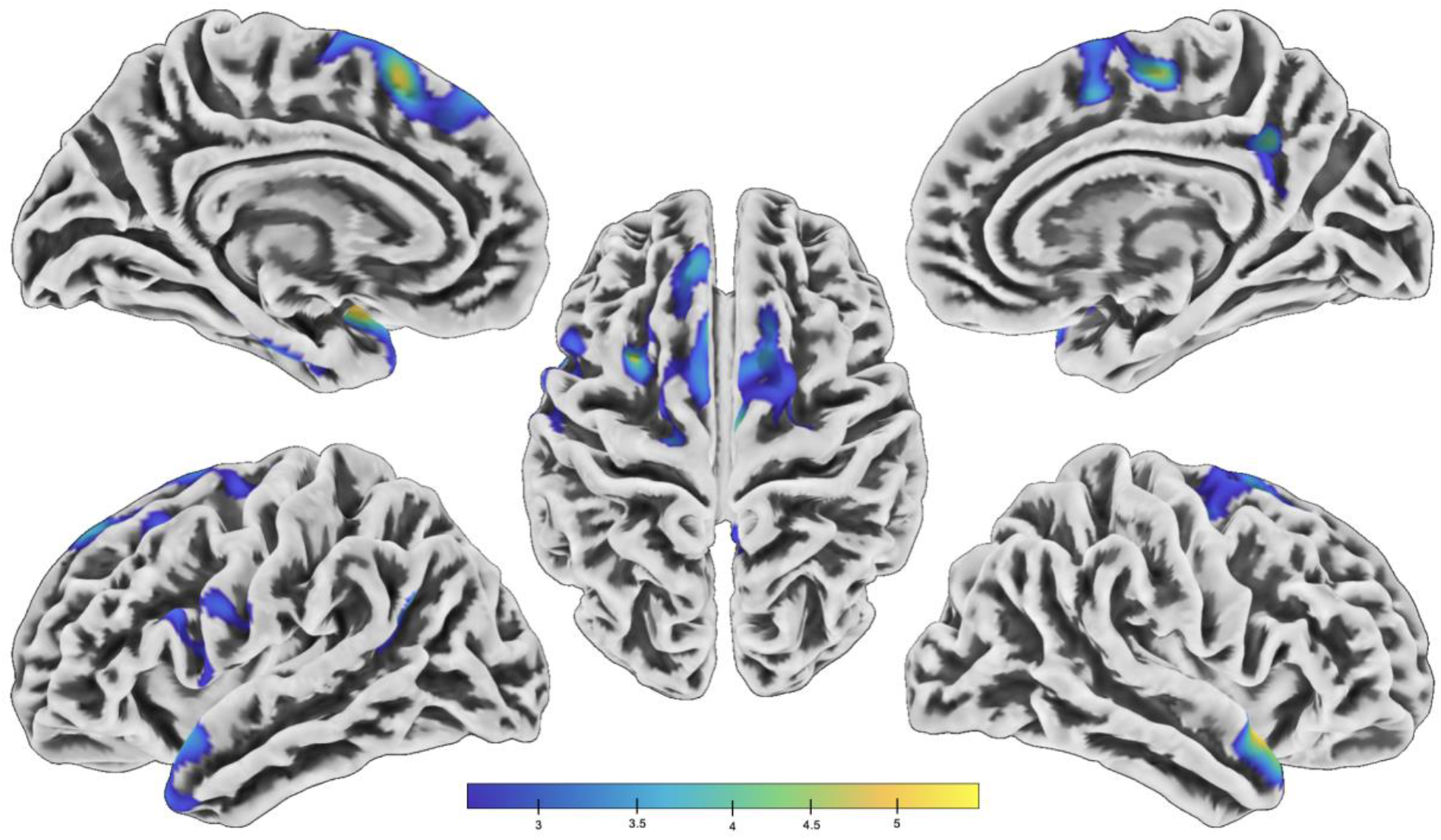
Cortical Thickness Results. Main Effect of Language Experience: Voxel-wise height threshold p < 0.005, cluster-level extent threshold p < 0.05, FDR. Coordinates for significant clusters for the main effect of Language Experience are provided in Table 5. (CT: n = 227)

#### CT: Main Effect of Diagnostic Group

There were no results from the main effect of Diagnostic Group on CT, indicating that the ADHD group did not differ from the non-ADHD Control group.

#### CT: Diagnostic Group x Language Experience Interaction

There were no results from the interaction between Diagnostic Group and Language Experience on CT.

## Discussion

The goal of the current study was to determine if an early dual-language experience has a modulating effect on the differences in EF performance and on the differences in neuroanatomy that manifest in ADHD. EF performance and structural brain differences in frontal-parietal and subcortical regions have been reported in both bilingualism and ADHD. However, these two lines of investigation have not converged to test whether the bilingual experience alters brain structure in these regions differentially in those with ADHD. Studies of ADHD have largely been conducted in monolinguals or participants whose language experience was not specified. Here we tested whether the two interact in ways to indicate that the neuroanatomical profile of ADHD is unique to bilinguals. Such a result would suggest that that findings from studies conducted in monolinguals with ADHD cannot be extrapolated to bilinguals with ADHD. We utilized a factorial design to compare early bilinguals and monolinguals with and without ADHD on three measures of EF performance, and on two neuroanatomical measures. We tested for an interaction between bilingualism and ADHD on EF performance and on neuroanatomy, with the expectation that a bilingual experience could stymie (or accentuate) the differences associated with ADHD. However, we found no interaction effects for any of the measures. As such, our results suggest that models of ADHD generated from studies conducted primarily in monolinguals are generalizable to bilinguals, and it is not likely that the bilingual experience is either beneficial or detrimental to ADHD.

### Behavioral Performance

For EF performance, the main effect of Language Experience yielded no evidence in support of better performance by bilinguals on any of the three tasks (Flanker, List Sort Working Memory, Dimensional Change Card Sorting). These results are surprising given some of the prior findings indicating a modification in bilinguals that leads to an ‘advantage’ in EF on similar measures of inhibitory control (Costa et al., 2009), working memory (Morales et al., 2013) and cognitive flexibility (Bialystok & Martin, 2004; Carlson & Meltzoff, 2008). As such, the results of the current study are more aligned with more recent research (Lehtonen et al., 2018; Lowe et al., 2021; Paap, 2019), including a large study using ABCD data (which involved the bilinguals in the current report as well as those using language pairs other than Spanish and English), which did not support the notion that EF is better in those who are bilingual relative to monolinguals (Dick et al., 2019).

For the main effect of Diagnostic Group, poorer performance for the ADHD Group compared to non-ADHD Control Group was only revealed on the Dimensional Change Card Sorting Task, but there were no effects on the other two tasks (Flanker and List Sort Working Memory Tasks). The absence of a group difference on the Flanker Task was unexpected given the well-established literature on inhibitory (interference) control deficits in ADHD (Mullane et al., 2009), and evidence that, on average, children with ADHD perform below their peers on measures of executive functioning more broadly (Barkley, 2006). Similarly, a null finding for the List Sort Working Memory Task diverges from prior work noting working memory deficits in ADHD (Brocki et al., 2008; Ramos et al., 2020). Other studies using ABCD Study data have found significant differences between children with ADHD and controls on the Flanker Task, but not consistently on the List Sort Working Memory Task, with effect sizes smaller than those reported in convenience samples and with high variability within the ADHD population (Cordova et al., 2022; Harkness et al., 2025; Sarabin et al., 2023). However, despite the theoretical consensus linking ADHD to EF deficits, a meta-analysis found that while children with ADHD showed statistically significant impairments across multiple EF domains, effect sizes were moderate, and no single EF deficit was present in all who have ADHD (Willcutt et al., 2005). Notably, a substantial proportion of children meeting diagnostic criteria for ADHD show no measurable EF impairment (Nigg et al., 2005; Willcutt et al., 2005), emphasizing that EF deficits in ADHD represent a symptom rather than a universal feature of the disorder.

Despite these null results in bilinguals and the weak results in those with ADHD on differences in EF performance, it is still possible that performance on these tasks could be modulated by the dual-language experience, but none of the ANOVAs on Flanker, List Sort Working Memory, or Dimensional Change Card Sorting Tasks yielded an interaction effect between Diagnostic Group (ADHD vs. Control) and Language Experience (Bilingual vs. Monolingual). Together these results indicate that a bilingual experience does not differentially affect executive function in those with ADHD.

### Neuroanatomical Differences Associated with Bilingualism

The main effect of language experience (ADHD and Control Groups combined) revealed extensive neuroanatomical differences between bilinguals and monolinguals, with bilinguals showing greater GMV across frontal, temporal, parietal, subcortical, and occipital regions. Many of these regions align with prior neuroanatomical studies of bilinguals (García-Pentón et al., 2016, 2019; Mechelli et al., 2004; Olulade et al., 2016; Schug et al., 2022; Schug & Eden, 2026). The pattern of more GMV in language regions, including the left inferior frontal gyrus, left supplementary motor area, bilateral temporal cortices, and thalamus is consistent with the cognitive demands of managing two languages, including speech production, semantic processing, and language (Mechelli et al., 2004; Pliatsikas, 2020; Schug et al., 2022; Nguyen et al., 2024). More GMV in attention and executive function regions, including right inferior frontal gyrus, bilateral precuneus, and left postcentral-superior parietal cortex extends these findings to domain-general networks (Olulade et al., 2016; Schug et al., 2022; Wang et al., 2025). On the other hand, bilinguals showed less GMV than monolinguals in posterior visual and cerebellar regions, suggesting differential specialization of these systems in the presence of dual-language experience. Together, these findings confirm prior work that dual-language experience affects cortical and subcortical regions broadly.

For CT, bilinguals had thinner cortex than monolinguals across frontal, temporal, and occipital regions of both hemispheres, with no regions showing relatively greater CT in bilinguals. These results were expected based on prior studies of bilingualism in the ABCD dataset, which included bilinguals of other language pairs (Vaughn et al., 2021). Cortical thinning in left-lateralized frontal regions, including the left supplementary motor area, middle frontal gyrus, inferior frontal gyrus, and precentral gyrus, suggests experience-dependent specialization of cognitive control and motor planning networks. Thinning in the superior temporal pole and left angular and fusiform gyri, is consistent with the demands of processing two languages. Thinner cortex in the right supplementary motor area and precuneus also indicates that the bilingual experience can shape cortical structure beyond language regions. Overall, neuroanatomical differences associated with bilingualism measured with GMV and CT are extensive, and notably more widespread than those associated with ADHD as described below.

### Neuroanatomical Differences Associated with ADHD

The main effect of diagnostic group (Bilingual and Monolingual Groups combined) revealed a small set of structural differences that align with the existing ADHD literature. For GMV, the group with ADHD showed more GMV than controls in the left superior orbitofrontal gyrus extending into the inferior orbitofrontal cortex, regions central to inhibitory control and decision-making. More GMV in ADHD in these regions may reflect delayed maturation, consistent with the maturational lag hypothesis of ADHD in which frontal regions show prolonged developmental trajectories relative to controls (Castellanos, 2002). On the other hand, there was less GMV in ADHD in the left thalamus and the middle occipital gyrus. Thalamic volume differences have been reported in ADHD, though findings are mixed. While a study (Ivanov et al., 2010) and meta-analysis (Nakao et al., 2011) reported reduced thalamic volume consistent with disruption of the cortico-striato-thalamo-cortical loop (Barkley, 2006), another reported greater thalamic volume, particularly in familial ADHD (Baboli et al., 2022) and in the inattentive subtype (Fu et al., 2021).

Turning to CT, we did not identify any differences. While this null finding diverges from prior studies reporting thinner cortex in ADHD, especially in prefrontal areas (Narr et al., 2009; Shaw et al., 2006; Silk et al., 2016), it is possible that, like earlier research into structural differences generally, these studies applied lower statistical thresholds and used smaller samples, as was typical in all fields at that time. Further, null results were not likely to have been published. Recent large-scale studies which afford greater statistical power and have employed more rigorous controls and conservative thresholds, have yielded more nuanced findings than the earlier work, with some reporting null CT findings. Notably, two prior studies of ADHD using ABCD Study data also yielded no between-group differences in CT (Bernanke et al., 2022; Sarabin et al., 2023), while another reported CT differences in few regions, including the insula and anterior cingulate cortex (Reimann et al., 2024). These results suggest that neuroanatomical differences in ADHD are less pronounced than previously believed (Pereira-Sanchez & Castellanos, 2021). While this does not eliminate the possibility that there may be regions that are differentially affected when ADHD occurs in the presence of bilingualism, we found there to be none, as discussed next.

### No Interaction effect between Bilingualism and ADHD on Neuroanatomy

As noted above, those with ADHD had *less* GMV than non-ADHD Controls (bilinguals and monolinguals combined) in left thalamus, while bilinguals (ADHD and non-ADHD Controls combined) exhibited *more* GMV in the thalamus than monolinguals. This confirms what was predicted from observations reported in the literature, that neuroanatomical aberrations associated with ADHD manifesting as less GMV reside in the same anatomical regions where the bilingual experience drives greater GMV. As noted in the introduction, we had expected to find this in frontal-parietal as well as basal ganglia structures, however there were none. Studies reporting less GMV in the thalamus in ADHD have already been discussed above, yet it is worth noting that several studies have reported more GMV in the thalamus in bilinguals and related these to language production (Burgaleta et al., 2016) and language selection (Nguyen et al., 2024). There was more GMV in the group with ADHD in left orbitofrontal gyrus, but this did not overlap with the location of any differences attributed to bilingualism (although there were some in nearby left superior and middle frontal gyri). Further, the left middle occipital gyrus had less GMV in ADHD than controls, while the bilinguals had less GMV than monolinguals in this same region, suggesting a compounding effect. However, for both the left thalamus and middle occipital gyrus, the degree of difference between those with ADHD and those without ADHD in the bilinguals versus monolinguals, was not sufficiently minimized or accentuated, respectively, to lead to an interaction effect (antagonistic or exponential). Importantly, the absence of a significant interaction in either GMV or CT alleviates concerns that a dual-language experience modulates neuroanatomy in children with ADHD and fits with the behavioral results on EF performance. From a theoretical standpoint this means that the neuroanatomical profile of ADHD does not appear to differ as a function of early bilingual language experience, suggesting that existing models developed primarily in monolinguals are applicable to bilingual children. From a practical standpoint, these results suggest that raising a child with ADHD in a bilingual environment is not likely to improve or worsen their ADHD, which should be reassuring for bilingual families and clinicians working with this population. By bringing together two fields that have largely operated in isolation, this work deepens our understanding of the extent and the limitations of the dual-language experience in shaping the brain.

## Conclusion

Disparate lines of research have investigated differences in ADHD and in bilinguals for executive function and brain structure. Here we provide the first investigation into bilinguals with ADHD to test if the performance and brain structural differences associated with ADHD are influenced by a dual-language experience. The results, based on 244 children from the ABCD Study, demonstrate that bilingualism is not associated with EF performance strengths, but does come with widespread neuroanatomical differences in GMV and CT, while ADHD is associated with some EF performance deficits and some cortical and subcortical differences in GMV, but not in CT. There was spatial convergence for the effects of bilingualism and ADHD in the left thalamus and middle occipital gyrus, but no interaction results to indicate that the difference between ADHD and non-ADHD controls is more pronounced if bilingualism is present. The results indicate that the framework for ADHD research and clinical practice established in monolinguals is also relevant to bilinguals.

## Supporting information

OakSupplement

## Data Availability Statement

All data files are available from the ABCD Study Data Repository from the NIMH Data Archive (https://nda.nih.gov/abcd).

## Acknowledgements

Data used in the preparation of this article were obtained from the Adolescent Brain Cognitive Development (ABCD) Study (https://abcdstudy.org), held in the NIMH Data Archive (NDA). This is a multisite, longitudinal study designed to recruit more than 10,000 children age 9–10 and follow them over 10 years into early adulthood. The ABCD Study® is supported by the National Institutes of Health and additional federal partners under award numbers U01DA041048, U01DA050989, U01DA051016, U01DA041022, U01DA0510 18, U01DA051037, U01DA050987, U01DA041174, U01DA041106, U01DA041117, U01DA041028, U01DA041134, U01DA050988, U01DA051039, U01DA041156, U01DA 041025, U01DA041120, U01DA051038, U01DA041148, U01DA041093, U01DA041089, U24DA041123, U24DA041147. A full list of supporters is available at https://abcdstudy.org/federal-partners.html.

The ABCD data repository grows and changes over time. The ABCD data used in this report came from the Fast Track data release. The raw data are available at https://nda.nih.gov/edit_collection.html?id=2573.

## Ethics Approval Statement

This study was performed in line with the principles of the Declaration of Helsinki. Parent consent and child assent was obtained by the ABCD Study (Garavan et al., 2018).

## Permission to reproduce material from other sources

N/A

