## Supplementary material for "Are executive function and neuroanatomy in ADHD modulated by bilingualism?": OakSupplement

**Supplementary Figure 1: Participant Selection Flowchart**

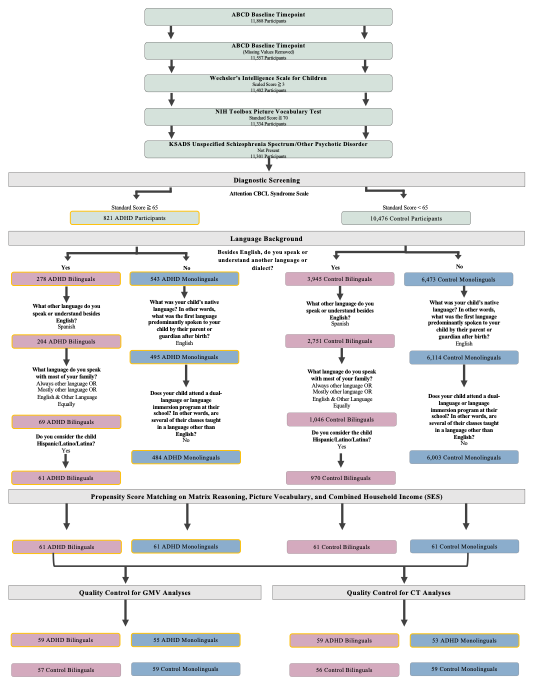

Overview of inclusion/exclusion criteria, propensity matching, and quality control from the ABCD Study sample leading to the final groups.

**Supplementary Table 1: Race Distribution for the Bilingual and Monolingual groups with and without ADHD.**

|  | **Bilingual ADHD**  (n=61) | **Bilingual Control** (n=61) | **Monolingual ADHD**  (n=61) | **Monolingual Control**  (n=61) |
| --- | --- | --- | --- | --- |
| White, Other Race | 4 | 0 | 0 | 0 |
| White | 37 | 33 | 29 | 28 |
| Black/African American | 2 | 1 | 19 | 21 |
| White, Black/African American | 2 | 1 | 4 | 3 |
| White, American Indian/Native American | 1 | 0 | 0 | 1 |
| White, American Indian/Native American, Other Race | 0 | 1 | 0 | 0 |
| Other Pacific Islander | 0 | 1 | 0 | 0 |
| White, Japanese | 0 | 0 | 1 | 0 |
| White, Black/African American, American Indian/Native American | 0 | 0 | 4 | 1 |
| White, Filipino/Filipina | 0 | 0 | 1 | 0 |
| Black/African American, Other Race | 0 | 0 | 1 | 0 |
| White, Korean | 0 | 0 | 1 | 0 |
| American Indian/Native American | 0 | 0 | 0 | 2 |
| White, Other Pacific Islander | 0 | 0 | 0 | 1 |
| White, Black/African American, American Indian/Native American, Other Pacific Islander | 0 | 0 | 0 | 1 |
| Other Race | 9 | 20 | 1 | 3 |
| Don't Know | 5 | 2 | 0 | 0 |
| Refuse To Answer | 1 | 2 | 0 | 0 |

*Note*: Group counts for racial identification as reported by a parent in response to the question "What race do you consider the child to be?".
